# Disparity sensitivity and binocular integration in mouse visual cortex areas

**DOI:** 10.1101/2020.05.07.083329

**Authors:** Alessandro La Chioma, Tobias Bonhoeffer, Mark Hübener

## Abstract

Binocular disparity, the difference between the two eyes’ images, is a powerful cue to generate the three-dimensional depth percept known as stereopsis. In primates, binocular disparity is processed in multiple areas of the visual cortex, with distinct contributions of higher areas to specific aspects of depth perception. Mice, too, can perceive stereoscopic depth, and neurons in primary visual cortex (V1) and higher-order, lateromedial (LM) and rostrolateral (RL) areas were found to be sensitive to binocular disparity. A detailed characterization of disparity tuning properties across mouse visual areas is lacking, however, and acquiring such data might help clarifying the role of higher areas for disparity processing and establishing putative functional correspondences to primate areas. We used two-photon calcium imaging to characterize the disparity tuning properties of neurons in mouse visual areas V1, LM, and RL in response to dichoptically presented binocular gratings, as well as correlated and anticorrelated random dot stereograms (RDS). In all three areas, many neurons were tuned to disparity, showing strong response facilitation or suppression at optimal or null disparity, respectively. This was even the case in neurons classified as monocular by conventional ocular dominance measurements. Spatial clustering of similarly tuned neurons was observed at a scale of about 10 μm. Finally, we probed neurons’ sensitivity to true stereo correspondence by comparing responses to correlated and anticorrelated RDS. Area LM, akin to primate ventral visual stream areas, showed higher selectivity for correlated stimuli and reduced anticorrelated responses, indicating higher-level disparity processing in LM compared to V1 and RL.

## Introduction

A fundamental ability of the mammalian visual system is combining information from both eyes into a unified percept of the three-dimensional world. Each eyes’ retina receives an image of the environment from a slightly different vantage point, such that a given object’s image can fall on non-corresponding locations on the two retinae, depending on the object’s distance from the observer. The accurate sensing of the difference between the two retinal images, called binocular disparity, is the first critical step underlying binocular fusion and stereoscopic depth perception (Gonzalez and Perez, 1998). In primates, many visual cortex areas play a role in this task, with distinct representations of binocular disparity among different areas (Parker, 2007; Welchman, 2016). For example, compared to the primary visual cortex (V1), neurons in extrastriate areas show broader disparity tuning and encode a wider range of disparities. Likewise, tuning curves of most V1 neurons are symmetric, but are often asymmetric in extrastriate areas (Gonzalez and Perez, 1998; Cumming and DeAngelis, 2001). Moreover, neurons in V1 and in dorsal stream areas, such as the middle temporal area (MT) and the medial superior temporal area (MST), respond to both binocularly correlated and anticorrelated stimuli (Cumming and Parker, 1997; Takemura et al., 2001; Krug, 2004), whereas ventral stream areas, such as V4 and inferior temporal cortex, display weaker or no responses to anticorrelated stimuli, reflecting higher-level processing of disparity signals and a close correlation with the perception of stereo depth (Janssen et al., 2003; Tanabe et al., 2004). Finally, visual areas across ventral and dorsal streams show different selectivities for either near or far stimuli (Cléry et al., 2018; Nasr and Tootell, 2018). Thus, the tuning characteristics of disparity sensitive neurons in the primate visual system have helped clarifying the hierarchy of visual areas and their distinct roles for stereo-based depth processing.

The mouse has become a key model for understanding visual cortex function, largely because of its experimental tractability (Niell, 2015; Glickfeld and Olsen, 2017). As in other mammals, mouse visual cortex consists of V1 and multiple higher areas with specific interconnections and different proposed roles for visual information processing, which are still relatively poorly understood, however (Wang et al., 2011, 2012; Andermann et al., 2011; Marshel et al., 2011; Zhuang et al., 2017; Han et al., 2018; de Vries et al., 2020).

Mice can discriminate stereoscopic depth (Samonds et al., 2019) and disparity-sensitive neurons similar to those characterized in other mammals have been found in V1 (Scholl et al., 2013) and in higher-order areas LM and RL (La Chioma et al., 2019). Clear differences in neurons’ preferred disparities were observed across these areas, with area RL being specialized for disparities corresponding to nearby visual stimuli (La Chioma et al., 2019).

Beyond these differences, binocular disparity has not been analyzed in detail in the different areas of mouse visual cortex. Obtaining this information should not only contribute to a better understanding of the role of higher areas for visual processing but it will also facilitate establishing functional correspondences to primate areas. Here, we characterize binocular disparity in V1 and areas LM and RL, which jointly contain the largest representation of the binocular visual field across mouse visual cortex. Using two-photon calcium imaging, we determined the disparity tuning properties of neurons using dichoptically presented binocular gratings, as well as correlated and anticorrelated random dot stereograms (RDS).

We found that, across these areas, many neurons were tuned to disparity. Binocularly presented stimuli caused strong response facilitation or suppression at optimal or null disparity, respectively, even in neurons classified as monocular by conventional ocular dominance measurements. While none of the areas studied showed a large-scale spatial organization for disparity preference, nearby neurons within 10 μm had similar tuning properties. Finally, a fraction of neurons across areas responded to anticorrelated RDS, with tuning curves inverted compared to correlated RDS. Area LM, similar to primate ventral visual stream areas, showed substantially fewer anticorrelated responses, suggesting a higher-level analysis of disparity signals in LM compared to V1 and RL.

## Results

To investigate binocular integration and disparity selectivity across mouse visual cortex, we performed *in vivo* two-photon calcium imaging in the primary visual cortex (V1), and areas LM and RL. We focused on these three areas since they contain the largest, continuous cortical representation of the central, binocular region of the visual field.

### Identification and targeting of areas V1, LM, and RL for two-photon imaging

To localize areas V1, LM, and RL for subsequent two-photon imaging, we used intrinsic signal imaging to map the overall retinotopic organization of the visual cortex (Fig. 1A; Kalatsky and Stryker, 2003; Marshel et al., 2011). Continuously drifting, horizontally and vertically moving bar stimuli were employed to generate two orthogonal retinotopic maps with precise vertical and horizontal meridians. By using the established visual field representations in mouse visual cortex (Marshel et al., 2011; Garrett et al., 2014), the boundaries among areas V1, LM, and RL could be readily identified (Fig. 1B,C).

**Figure 1.**
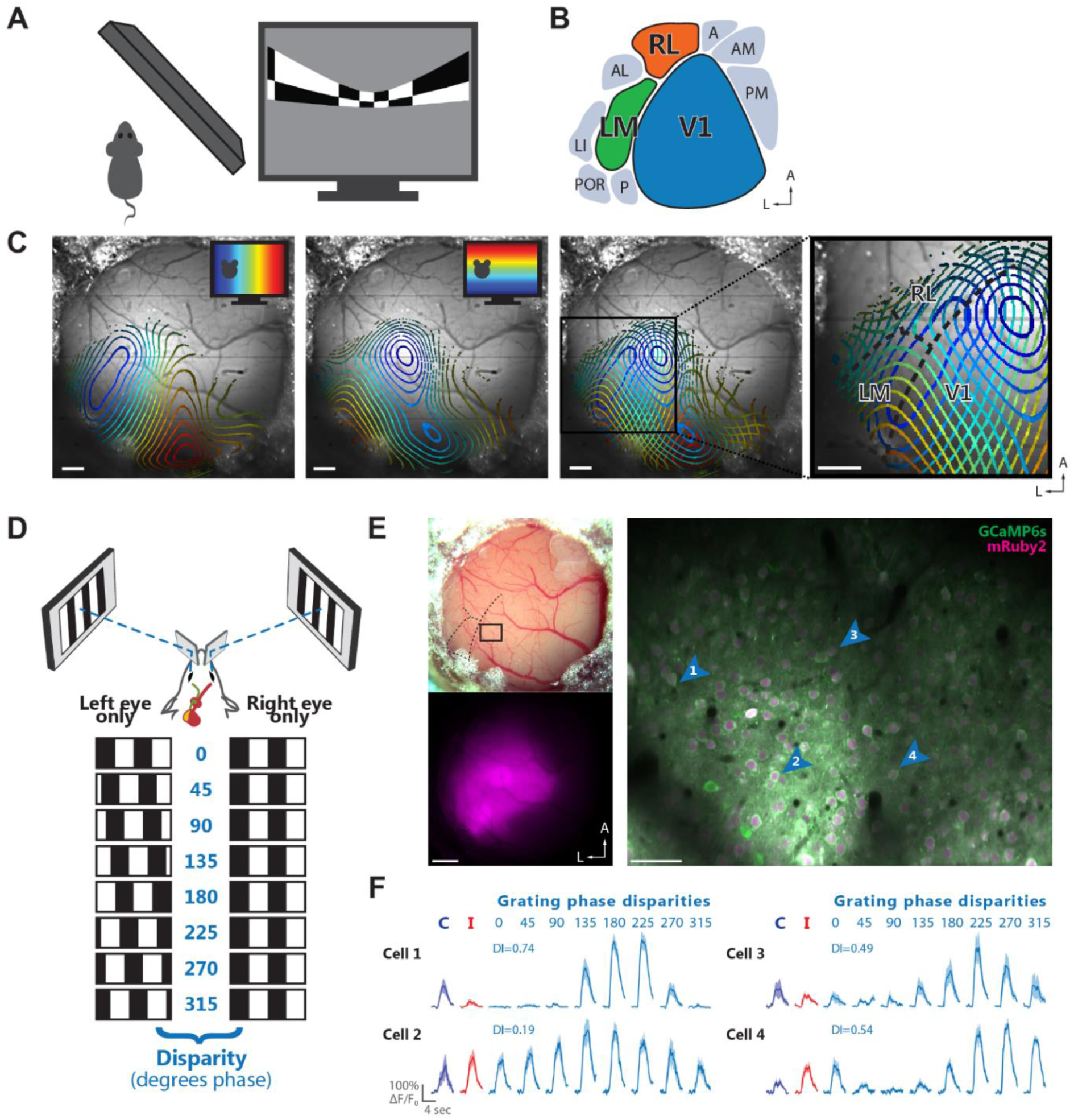
Identification and targeting of areas V1, LM, and RL for two-photon calcium imaging. (A) Schematic of stimulus presentation for mapping the retinotopic organization of mouse visual cortex areas. Left, top view; right, periodic bar stimulus displayed with spherical correction. (B) Schematic of the location of V1 and several higher-order areas of mouse visual cortex in the left hemisphere. The color code for areas V1 (blue), LM (green), and RL (orange) is used throughout the Figures. (C) Retinotopic maps from an example mouse. Contour plots of retinotopy are overlaid with an image of the brain surface. Contour lines depict equally spaced, iso-elevation and iso-azimuth lines as indicated by the color code. Panels from left to right: contour plot for azimuth; contour plot for elevation; overlay of azimuth and elevation contours; enlarged view of cortical areas V1, LM, and RL. The boundaries between these areas (dashed black lines) can be reliably delineated. Scale bars, 500 μm. (D) Schematic illustrating dichoptic grating stimulation. Top, haploscope apparatus for dichoptic presentation of visual stimuli. Bottom, drifting gratings are dichoptically presented at varying interocular phase disparities. Eight equally spaced interocular grating disparities (0–315 deg phase) are produced by systematically varying the initial phase (position) of the grating presented to one eye relative to the phase of the grating presented to the other eye. (E) Two-photon imaging using the calcium indicator GCaMP6s co-expressed with the structural marker mRuby2. Top left, image of a cranial window 6.5 weeks after implantation. Bottom left, epifluorescence image showing the expression bolus, with fluorescence signal from mRuby2. Right, example two-photon imaging plane acquired ∼180 μm below the cortical surface in area V1. The image shows a mean-intensity projection (20,000 frames, shift corrected) with fluorescence signal from GCaMP6s (green) and mRuby2 (magenta). The cortical location of this imaging plane is indicated in the top left panel. Scale bars: left panels, 500 μm; right panel, 50 μm. (F) Visually-evoked calcium transients (ΔF/F_0_) of four example neurons indicated by the blue arrowheads in (E). For each cell, the responses to monocular drifting gratings presented to either the contralateral (blue) or ipsilateral eye (red) are shown on the left. Responses to the eight interocular phase disparities of dichoptic gratings are shown on the right (cyan), along with the corresponding disparity selectivity index (DI). The fluorescence time courses are plotted as mean ΔF/F_0_ and SEM (lines and shaded areas) calculated across stimulus trials.

After having identified V1, LM, and RL, we targeted these areas for two-photon imaging (Fig. 1E,F). Visually-evoked activity of individual neurons was measured using the genetically encoded calcium indicator GCaMP6s (Chen et al., 2013), co-expressed with the structural marker mRuby2 (Rose et al., 2016). The latter aided image registration for correcting motion artifacts and improved the identification of neurons for marking ROIs.

### Binocular disparity is encoded by large fractions of neurons in areas V1, LM, and RL

Disparity tuning was characterized using drifting vertical gratings displayed in a dichoptic fashion at varying interocular disparities (Fig. 1D,F). Eight different grating disparities were generated by systematically varying the relative phase between the gratings presented to either eye, while drift direction, speed, and spatial frequency were kept constant across eyes. In addition to such “dichoptic gratings”, gratings were also displayed to each eye separately (“monocular gratings”) to determine ocular dominance (OD) and compare monocular with binocular responses.

To minimize the “correspondence problem” that arises when using circularly repetitive stimuli, dichoptic grating stimuli were presented at a low spatial frequency (0.01 cycles per degree, i.e., 100 degrees per cycle), such that for most cells no more than a single grating cycle is covered by individual receptive fields (RFs), given the typical RF size in mouse visual areas (Van den Bergh et al., 2010; Smith et al., 2017) and a binocular overlap of about 40 degrees (Scholl et al., 2013; Sterratt et al., 2013).

Across areas, ∼15% of neurons were responsive (see Materials and Methods) to vertical, dichoptic gratings (percentage of responsive cells, mean ± SEM across planes, V1, 14.1% ± 2.0%; LM, 15.8% ± 1.4%; RL, 19.0% ± 0.9%). Altogether, the mean response magnitude was similar across areas (ΔF/F, mean ± SEM across planes, V1, 57% ± 3%; LM, 65% ± 6%; RL, 55% ± 3%).

For each responsive cell, a disparity tuning curve was computed by plotting its average calcium response in function of the interocular disparity of the dichoptic gratings (Fig. 2A). Typically, across areas, disparity tuning curves of neurons showed a strong modulation. To quantify the magnitude of modulation caused by binocular disparity, a disparity selectivity index (DI), based on the vectorial sum of responses across disparities (Scholl et al., 2013; La Chioma et al., 2019), was calculated for each cell, with values closer to one for highly selective cells and values closer to zero for less selective cells. Cells with DI > 0.3 were defined as disparity-tuned.

**Figure 2.**
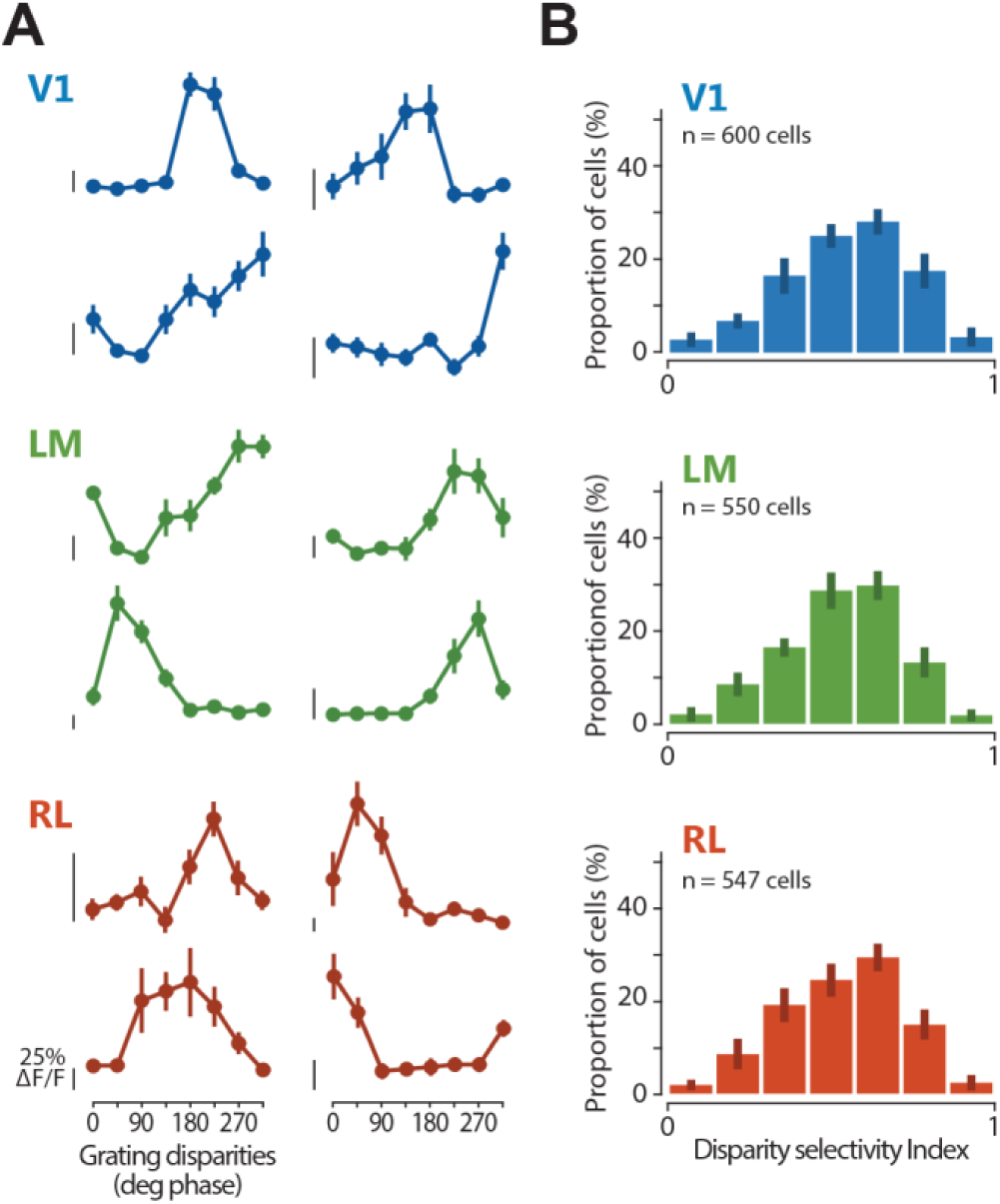
Functional characterization of disparity-tuned neurons in areas V1, LM, and RL. (A) Example tuning curves for binocular disparity, each from a different cell located in one of areas V1, LM, and RL as indicated by the color code. The mean fluorescence response is plotted as a function of the eight grating disparities. Error bars indicate SEM across trials. Scale bars for neuronal response indicate 25% ΔF/F, with the bottom end of each scale bar indicating the baseline level (0% ΔF/F). (B) Distributions of disparity selectivity index (DI) for each area. Mean of medians across planes ± SEM, V1, 0.57 ± 0.03; LM, 0.54 ± 0.02; RL, 0.55 ± 0.01.

To determine the overall sensitivity for binocular disparity across the three areas, the distribution of DI values of all cells was plotted (Fig. 2B). In all areas, the majority of neurons showed at least some degree of modulation to binocular disparity (percentage of responsive cells defined disparity-tuned, mean ± SEM across planes, V1, 89.0% ± 1.3%: LM, 88.4% ± 2.7%; RL, 88.3% ± 2.1%). Using more stringent criteria for defining responsive cells did not qualitatively change the DI distribution for each area (data not shown). Across all three areas, we observed overall similar degrees of disparity selectivity (Fig. 2B; Kruskal-Wallis test across planes, Χ^2^(2) = 1.9773, p = 0.3721; V1, n = 8 imaging planes, 600 responsive cells total, 7 mice; LM, n = 7 imaging planes, 550 responsive cells total, 7 mice; RL, n = 6 imaging planes, 547 responsive cells total, 5 mice). In addition, all three areas showed continuous DI distributions, indicating a continuum of disparity tuning, without pointing to the presence of a distinct subset of highly tuned cells. Thus, disparity sensitivity to gratings is widespread in V1 and higher areas LM and RL.

### Ocular dominance is similar across visual areas and is not correlated with disparity selectivity

By definition, disparity-tuned neurons are binocular. However, conventionally, a neuron’s binocularity is assessed by measuring its OD based on responses to monocular stimuli only. How are disparity selectivity and OD, which represent two different ways of describing a neuron’s binocularity, related to each other? To address this question, we first computed the ocular dominance index (ODI, ranging from +1 or −1) using the neuronal responses to monocular gratings presented to each eye separately. As reported (Dräger, 1975; Gordon and Stryker, 1996; Mrsic-Flogel et al., 2007; Rose et al., 2016), the distribution of ODI values for mouse V1 is considerably biased toward the contralateral eye (Fig. 3A). Neurons in areas LM and RL, for which OD measurements have not been reported yet, were also more strongly driven by the contralateral eye and showed ODI distributions comparable to V1 (ODI median, V1: 0.40, LM: 0.40, RL: 0.44; ODI mean ± SEM across experiments, V1: 0.17 ± 0.07, LM: 0.25 ± 0.06, RL: 0.21 ± 0.09; Kruskal-Wallis test for medians across planes Χ^2^(2) = 0.84, p = 0.658).

**Figure 3.**
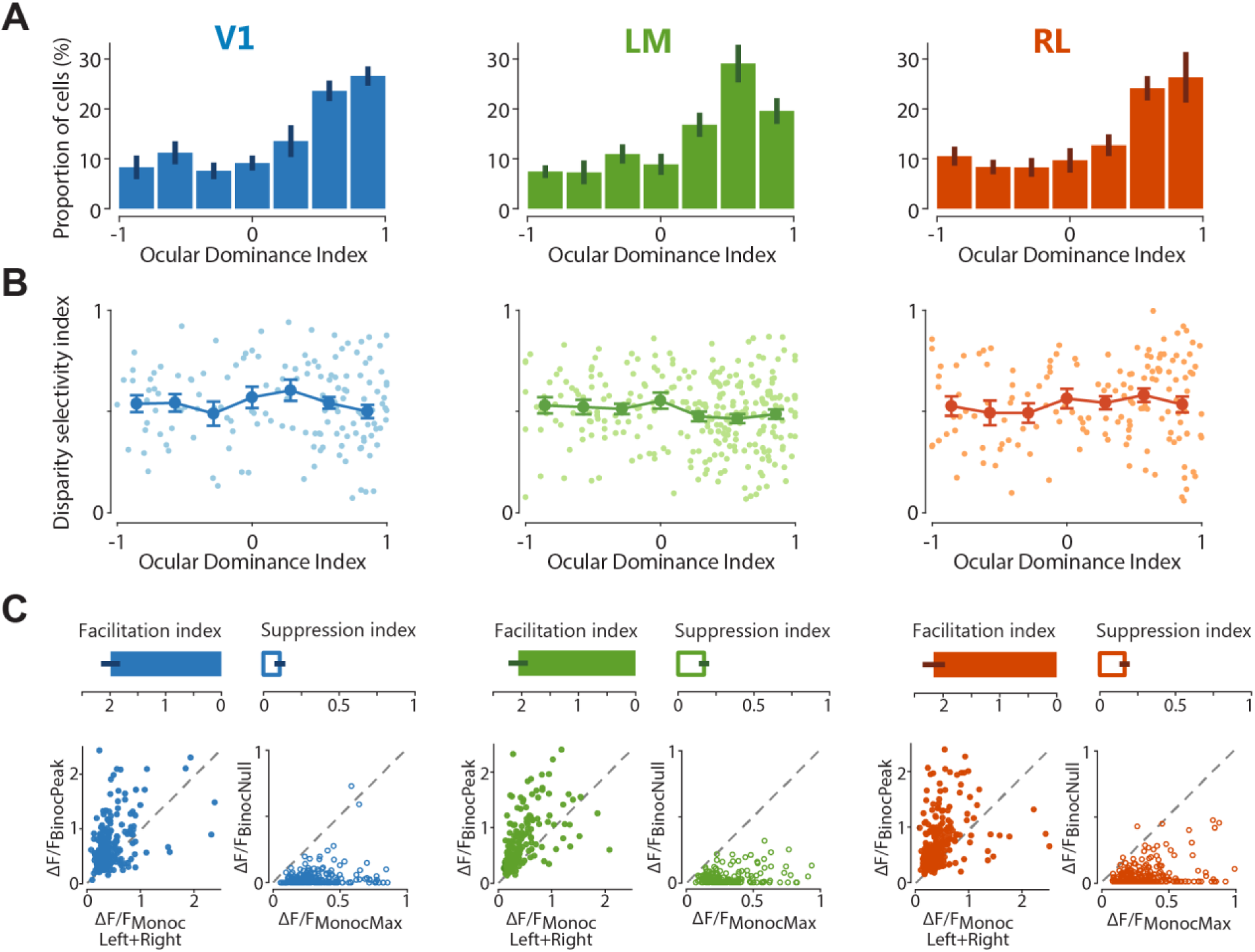
Ocular dominance and binocular interaction of individual neurons across visual areas. (A) Ocular dominance index (ODI) distributions in areas V1, LM, and RL, with bar plots indicating mean ± SEM across imaging planes. (B) Scatter graphs plotting the relationship between ODI and DI. Lines with error bars plot the mean ± SEM across neurons, with individual cells represented by dots in lighter shading. (C) Facilitatory and suppressive interactions upon binocular stimulation. Top panels, bar plots of facilitation and suppression indices for each area, with mean ± SEM across imaging planes. Bottom panels, for each area, the graph on the left plots the strongest response to dichoptic (binocular) gratings (ΔF/F_BinocPeak_) against the sum of the strongest contralateral and ipsilateral responses (ΔF/F_MonocLeft+Right_) for individual neurons, showing a strong overall response facilitation with binocular stimulation at the preferred disparity. For each area, the graph on the right plots the weakest response to dichoptic (binocular) gratings (ΔF/F_BinocNull_) against the strongest monocular response (ΔF/F_MonocMax_) for individual neurons, showing a strong overall response suppression with binocular stimulation at the least preferred (null) disparity.

We then analyzed the relationship between OD and disparity selectivity by plotting DI values against ODI values for individual cells in each area. Disparity-tuned neurons homogeneously covered the entire range of ODI values (Fig. 3B; Pearson’s correlation, V1: r = −0.03, p = 0.706; LM: r = 0.07, p = 0.390; RL: r = −0.10, p = 0.095; Kruskal-Wallis test across ODI bins: V1, Χ^2^(6) = 4.5563, p = 0.6018; LM, Χ^2^(6) = 8.1326, p = 0.2286; RL, Χ^2^(6) = 3.0209, p = 0.8062). Notably, neurons classified as monocular by OD measurements (ODI ≈ 1 or ODI ≈ −1) could be disparity-tuned, hence clearly reflecting integration of inputs from both eyes. Thus, there is no clear relationship between OD and disparity selectivity, in line with other studies in mice (Scholl et al., 2013), cats (Ohzawa and Freeman, 1986; LeVay and Voigt, 1988), and monkeys (Prince et al., 2002; Read and Cumming, 2003).

### Most neurons exhibit strong disparity-dependent facilitation and suppression

To examine the integration of visual inputs from both eyes, we next compared the responses evoked by dichoptic and monocular gratings. The response at the optimal binocular disparity was generally much larger than the sum of the two monocular responses for most neurons, indicating a robust facilitatory effect of binocular integration (Fig. 3C). At the same time, there were also strong suppressive binocular interactions: at the least effective disparity, the response was generally absent or smaller than the larger of the two monocular responses (Fig. 3C).

To quantify the response facilitation or suppression at the preferred and least-preferred disparity, respectively, a facilitation index (FI) and a suppression index (SI) was computed for every neuron responsive to both dichoptic and monocular gratings (see Materials and Methods). FI values above 1 indicate a facilitatory interaction of the eye-specific inputs, while SI values below 1 correspond to suppressive binocular interactions. Most neurons across the three areas were overall highly and similarly susceptible to binocular interactions, as inputs from both eyes were integrated with strong response facilitation as well as suppression as a function of binocular disparity (Fig. 3C; one-way ANOVA for FI across areas: F(2,18) = 0.4353, p = 0.6537; SI, F(2,18) = 1.3776, p = 0.2775).

### Spatial clustering of neurons with similar disparity preference

While no large-scale spatial arrangement for response properties has been observed in mouse visual cortex (Mrsic-Flogel et al., 2007; Zariwala et al., 2011; Montijn et al., 2014), several studies have reported a fine-scale organization for orientation preference and OD (Ringach et al., 2016; Kondo et al., 2016; Maruoka et al., 2017; Scholl et al., 2017). To test whether binocular disparity preference is organized in a similar fashion, we generated color-coded, disparity maps for each imaging plane, with a cell body’s hue coding for its disparity preference (Fig. 4A). As expected, inspection of these maps did not reveal a large-scale arrangement of disparity-tuned neurons.

**Figure 4.**
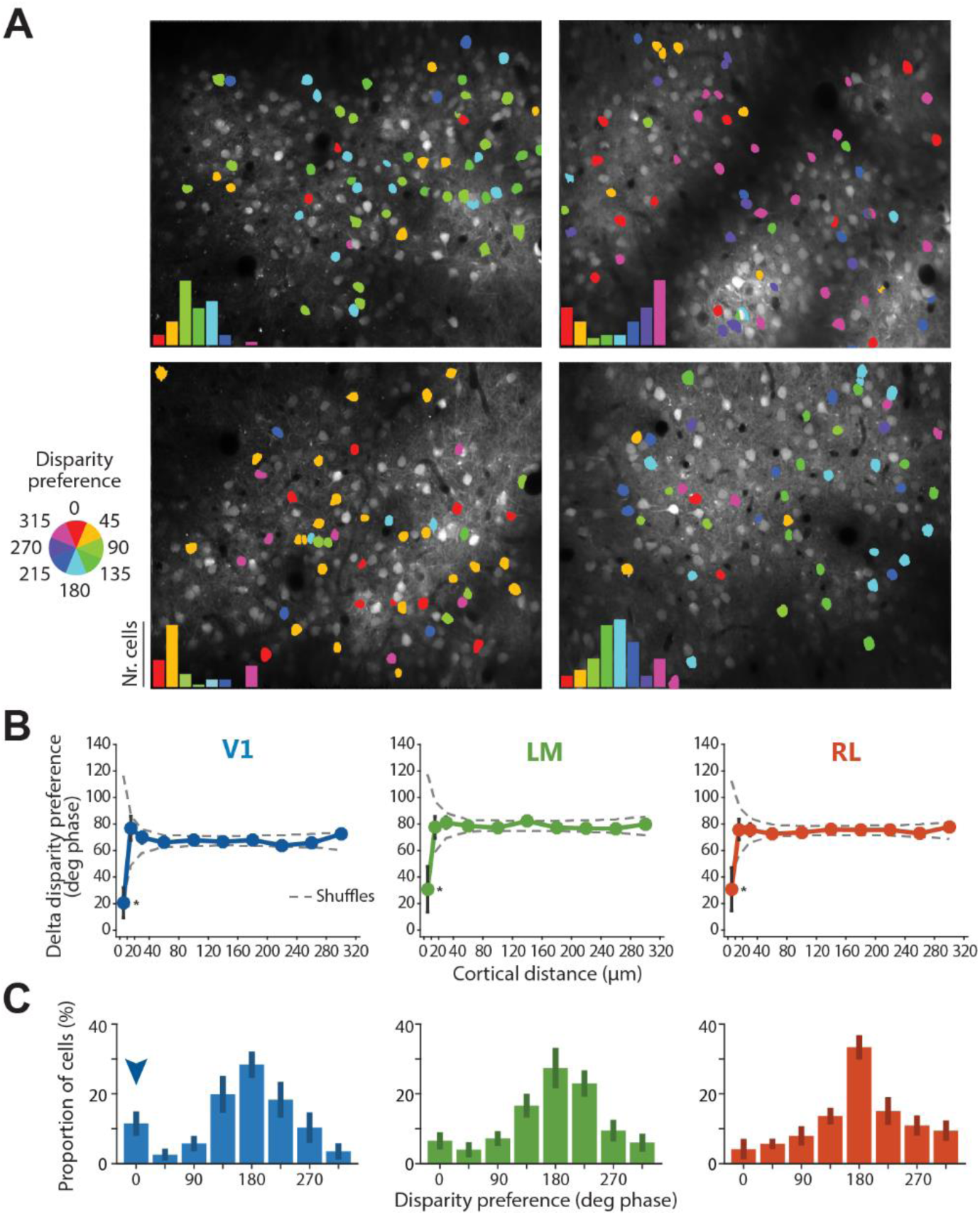
Spatial and functional organization of disparity-tuned neurons. (A) Example disparity maps from four different imaging planes, with neurons color-coded for disparity preference. Insets, disparity preferences show non-uniform distributions, with a population peak disparity characteristic for each individual experiment. (B) Spatial organization for disparity tuning. Difference in disparity preference between every pair of disparity-tuned cells in each imaging plane plotted as a function of the cortical distance between cells. Plot lines and error bars indicate mean ± SEM across neurons. The dashed lines indicate the 95% confidence interval, as determined by random shuffles of disparity preferences and cell x,y positions in each imaging plane (*p < 0.05 at the 10 μm cortical distance bin). (C) Peak aligned distributions of disparity preferences, averaged across experiments. The population peak was arbitrarily set to 180 deg phase. The arrowhead in the left panel for area V1 indicates a secondary peak of cells with disparity preference at 0 deg phase.

To investigate whether a spatial organization exists on a finer scale, we plotted the difference in disparity preference between every pair of cells as a function of their cortical distance (Fig. 4B). None of the three areas showed a clear dependence of tuning similarity on cortical distance, indicating a lack of a large-scale spatial arrangement of disparity-tuned cells. Nonetheless, adjacent neurons, located within 10 μm from each other, showed a preference for similar disparities, indicating some degree of spatial clustering on the scale of 10 μm. This spatial scale is consistent with values reported for orientation and spatial frequency tuning in mouse visual cortex (on the scale of ∼35 μm; Ringach et al., 2016; Scholl et al., 2017), and also consistent with reports demonstrating a fine-scale organization for orientation tuning and OD (in the range of 5–20 μm), with neurons sharing functional properties arranged into microcolumns (Kondo et al., 2016; Maruoka et al., 2017).

### Non-uniform distribution of disparity preference in individual experiments

While there was no obvious, large-scale spatial organization for disparity preference, in each individual experiment we found that disparity preferences showed a non-uniform distribution, with a peak characteristic for each experiment (Fig. 4A). This peak disparity varied over the whole range from experiment to experiment, showing no systematic relationship across experiments or animals. We consider it unlikely that the variation of apparent population peak disparity across mice reflects a true biological phenomenon. Most likely, these variations across experiments reflect differences in the alignment of the optical axes of the animal’s eyes, created mostly by a technical factor: namely the precise positioning of the haploscope apparatus, which could not be controlled with the necessary precision from experiment to experiment. It follows that the disparity preference distribution in each experiment is likely related to the actual optical axes of the mouse eyes in that imaging session, with the population peak disparity being approximately aligned with the visual field location of retinal correspondence between eyes (La Chioma et al. 2019). To determine the overall range of binocular disparity preferences in each area, the disparity preference distributions of individual experiments were aligned by setting the population peak arbitrarily to 180 deg phase and averaging the distributions across experiments (Fig. 4C). We observed that the disparity preference distributions were comparable across areas, apart from a higher fraction of cells with a disparity preference of 0 deg phase in area V1 (see arrowhead in Fig. 4C).

### Subsets of neurons in areas V1 and LM have asymmetric disparity tuning curves

The secondary peak in the distribution of disparity preferences in V1 points towards a subset of neurons with distinct properties. Two possible mechanisms have been put forward to explain the disparity selectivity of cells in the visual cortex. (1) According to the position-shift model, the disparity tuning of a neuron arises from a spatial offset in the RF position between left and right eye, with the RF of each eye having the same spatial arrangement of subfields. (2) Alternatively, the phase-shift model posits that a disparity-tuned neuron has left and right eye RFs at the same retinal position, but with different subfield structures. The position-shift model produces symmetric disparity tuning curves, whereas the phase-shift model is expected to cause asymmetric (or odd-symmetric) tuning curves (Qian, 1997; Cumming and DeAngelis, 2001; Tsao et al., 2003). We thus analyzed the tuning curves in more detail. First, we plotted the normalized tuning curves averaged across all disparity-tuned neurons from each area (Fig. 5A). Overall, the three areas showed very similar tuning curves. Next, we fitted the disparity tuning curves of individual neurons with an asymmetric Gaussian function with separate width parameters for the left (σ1) and right side (σ2; Fig. 5B; Hinkle and Connor, 2005; see Materials and Methods for details), allowing to assess potential tuning curve asymmetries. Indeed, asymmetry varied among cells, and systematically depended on the cells’ disparity preference (Fig. 5D): in V1 and LM, neurons with 0 deg phase disparity preference had asymmetric tuning curves, which were consistently skewed to the right (Fig. 5C,D). In contrast, neurons with a disparity preference close to the central peak of the distributions (at 180 deg phase) showed symmetric tuning curves (Fig. 5C,D). Therefore, these data suggest that the disparity selectivity of most neurons across mouse visual areas is mainly generated by a position-shift mechanism, whereas the phase-shift mechanism would be responsible for the disparity selectivity of a specific subset of neurons located mostly in V1 and LM.

**Figure 5.**
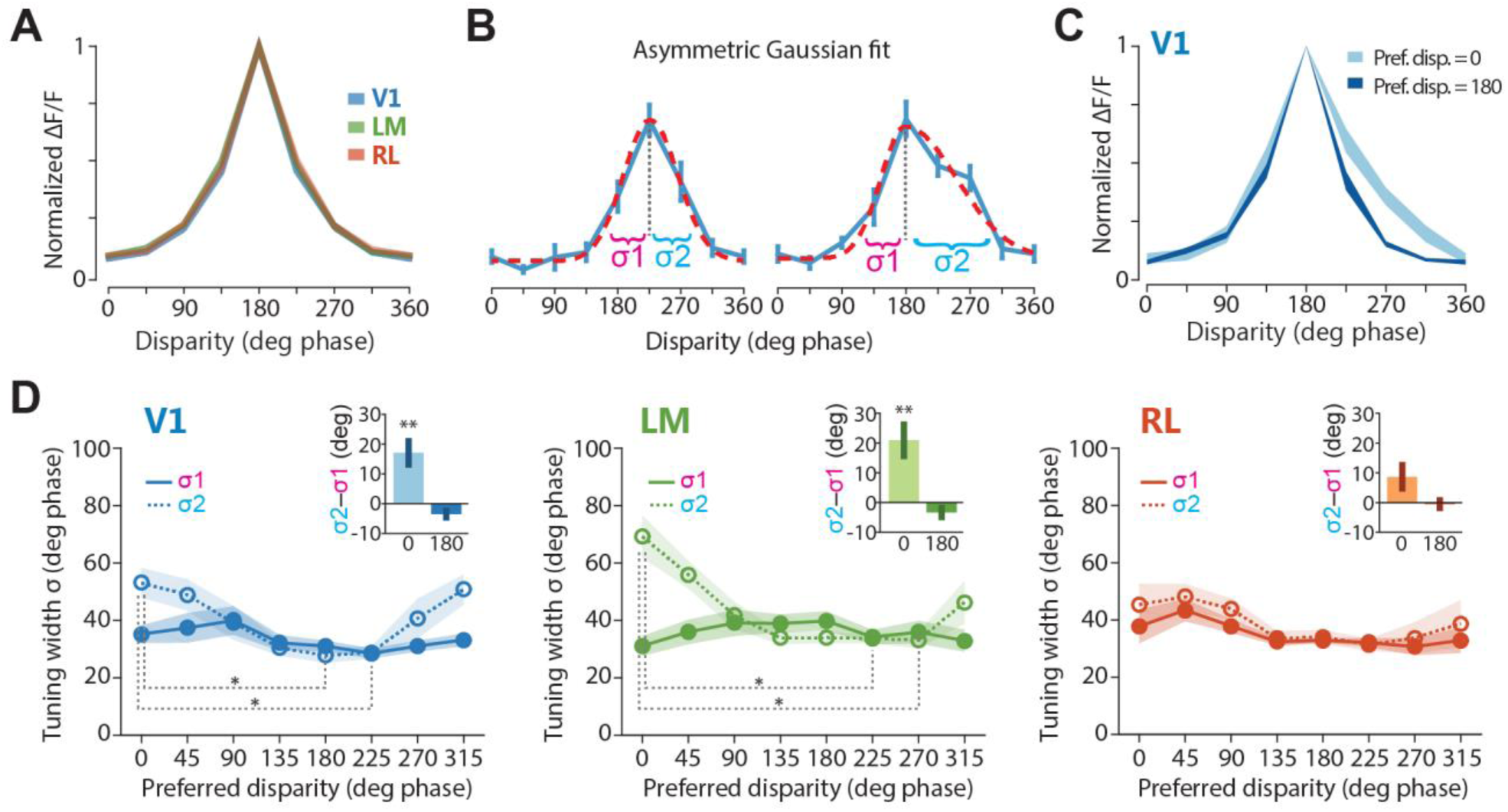
Analysis of disparity tuning curves. (A) Disparity tuning curves averaged across disparity-tuned neurons for each area. For averaging, each individual tuning curve was peak aligned to 180 deg phase and the response amplitudes normalized to the peak response. Shaded regions are ± SEM calculated across disparity-tuned neurons (DI > 0.3). (B) Tuning curve fit with an asymmetric Gaussian function. Two examples curve fits are shown, illustrating the tuning width parameters for the left and right sides (σ1 and σ2). The asymmetry of a tuning curve is quantified as the difference between the two width parameters (σ2 - σ1). Left, example of a symmetric tuning curve. Right, example of an asymmetric tuning curve. (C) Disparity tuning curves averaged across disparity-tuned neurons in V1, with separate averaging for neurons with disparity preference of 180 deg phase (dark blue) and 0 deg phase (light blue). Shaded regions are ± SEM. Disparity-tuned neurons with a disparity preference of 0 deg phase show tuning curves skewed to the right side of the peak. (D) Tuning width parameters plotted as a function of the disparity preference of each disparity-tuned cell. Lines with shaded regions indicate mean ± SEM calculated across imaging planes for the left (solid line) and right (dotted line) tuning width parameter. One-way ANOVA followed by multiple comparisons with Bonferroni correction. Insets, difference between right and left width parameters (σ2 - σ1) for neurons with disparity preference of 0 deg phase or 180 deg phase. Bar plots indicate mean ± SEM calculated across neurons. One sample t-test against zero.

### Noise correlations are higher between neurons with similar disparity preference

We next analyzed the trial-to-trial fluctuations in response strength between pairs of neurons, the so-called noise correlations. The amount of variability shared between neurons has implications for neural coding, and is assumed to reflect connectivity among cells (Averbeck et al., 2006; Cohen and Kohn, 2011; Ko et al., 2011; Schulz et al., 2015; Kohn et al., 2016), with highly correlated neurons being more strongly interconnected or sharing more common inputs compared to neurons with lower noise correlations (Ko et al., 2011). Over the entire populations, in each area, the pairwise noise correlations were on average weak but significantly larger than zero (Fig. 6A, insets; right-tailed t-test for mean > 0, V1, p = 0.0024, LM, p = 0.0009, RL, p = 0.0014), in line with previous reports in mouse visual cortex (Ko et al., 2011; Montijn et al., 2014; Rose et al., 2016; Khan et al., 2018). Moreover, the distributions of noise correlations were comparable across areas (one-way ANOVA, F(2,18) = 1.0054, p = 0.3855), and all showed a positive tail consisting of small numbers of highly correlated pairs (Fig. 6A).

**Figure 6.**
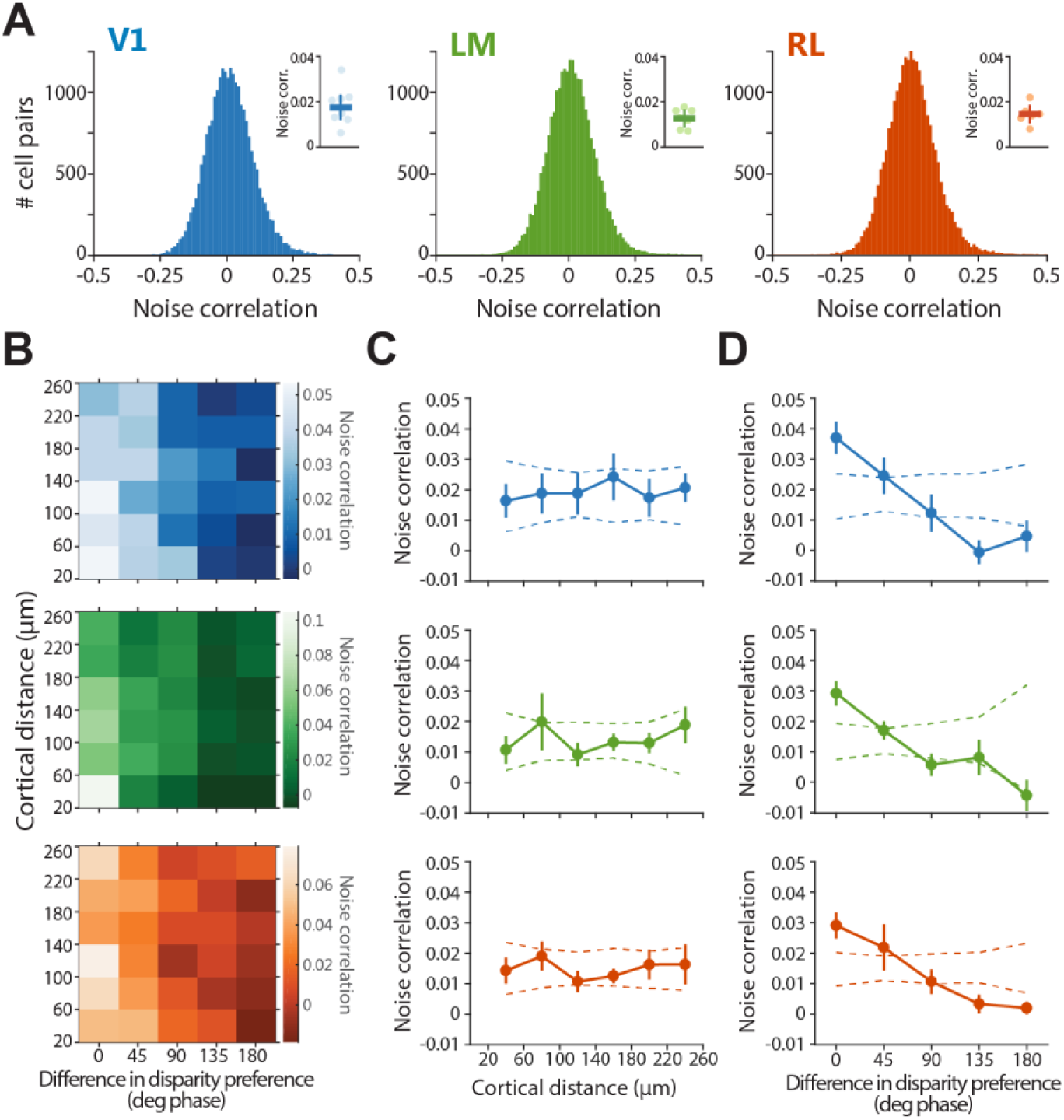
Noise correlations are higher between neurons with similar disparity preference. (A) Distributions of pairwise noise correlations. Only one distribution for each area is shown as an example. Note the small positive tail in the distributions. Insets, pairwise noise correlations averaged across planes ± SEM, with individual planes indicated with circles in lighter shading. (B) Dependence of noise correlations on both cortical distance and difference in disparity preference between each cell pair. (C) Pairwise noise correlation as a function of cortical distance between each cell pair. (D) Pairwise noise correlation as a function of difference in disparity preference between each cell pair. In (C-D), plot lot lines and error bars indicate mean ± SEM across imaging planes, and dashed lines indicate the 95% confidence interval, as determined by random shuffles of cell x,y positions in each imaging plane. For computing pairwise noise correlations, only cell pairs separated by at least 20 μm were considered.

It has been previously shown that the connection strengths between neurons in mouse visual cortex is stronger for cells with similar RFs and orientation tuning (Cossell et al., 2015), or for cells with high noise correlations (Ko et al., 2011), but it is not influenced by cortical distance (Cossell et al., 2015). Do nearby neurons or similarly disparity-tuned neurons have higher noise correlations? To answer this question, for each pair of disparity-tuned neurons, the value of their noise correlation was related to both their cortical distance and their disparity preference (Fig. 6B–D). Nearby and distal neurons showed comparable noise correlations, indicating no dependence of noise correlations on cortical distance (Fig. 6C). In contrast, pairs with a similar disparity preference (< 45 deg phase) showed substantially higher noise correlations compared to pairs with dissimilar preference (Fig. 6D). Thus, neurons with similar disparity preference are more strongly interconnected or share common inputs.

### Neuronal populations across visual areas effectively discriminate between grating disparities

Accurate representations of binocular disparity are likely encoded at the population level, since individual neurons are insufficient for this task, considering the narrow range of their response properties (Scholl et al., 2013; Burge and Geisler, 2014; Kato et al., 2016). Having shown that large numbers of individual neurons in all three areas encode binocular disparity, we next investigated how much information is carried, in each area, by the joint activity of multiple neurons.

We therefore employed a population decoding approach based on support vector machines (SVM; Cortes and Vapnik, 1995), trained using the calcium transients of populations of neurons. For each area, the SVM decoders were used to estimate, on a trial-to-trial basis, which of the eight grating disparities was actually presented (see Materials and Methods). Across areas, decoders were able to effectively estimate binocular disparity, since populations with as few as two neurons allowed significantly correct prediction of stimulus disparity, with initially steep improvement with increasing population sizes (Fig. 7). The three areas showed a similar capacity of discriminating disparity over the entire range of population sizes tested (Fig. 7). Moreover, a decoder built on a pseudo-population consisting of neurons pooled together from all three areas indistinctly, showed a curve of discrimination accuracy comparable to decoders trained on populations from each area separately (data not shown), thereby indicating that neurons from the different areas were interchangeable from the perspective of the decoder and hence contributed similarly to decoding. Together, these data indicate that populations of neurons in areas V1, LM, and RL efficiently encode binocular disparity and can effectively discriminate between grating disparities, with a comparable accuracy across areas.

**Figure 7.**
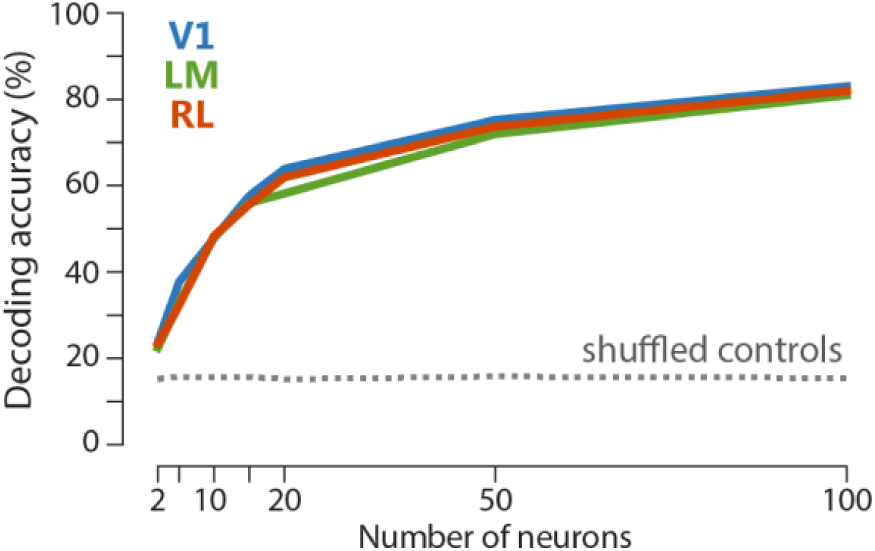
Population decoding of binocular disparity. Accuracy of linear SVM decoders trained to estimate which grating disparity, among all eight possible disparities, was actually presented. The classification accuracy of linear SVM decoders are plotted as a function of the number of neurons used for training the decoders, with neurons from each area (see color code). Plot lines indicate the mean accuracy across decoding iterations. Dashed line indicates the 95% confidence level, calculated as the 95^th^ percentile of the decoding accuracy after shuffling stimulus identity labels. See Materials and Methods for details.

### Area-specific responses to anticorrelated random dot stereograms

To extract depth information using binocular disparity, the visual system must determine which points in the left- and right-eye images correspond to the same visual feature among a number of potential false matches, i.e. the “correspondence problem” must be solved. Neurons sensitive to true stereo-correspondence should respond only to correct, but not false matches. Any sensitivity to false matches would indicate that further processing is needed for achieving true stereo-correspondence, e.g. in downstream areas. In primates, comparing the sensitivity to true and false matches across visual areas helped delineating their hierarchy and role for stereo-based depth processing (Parker, 2007). To test this in mouse visual cortex, we probed disparity tuning in awake animals using RDS, comparing responses to binocularly correlated RDS (cRDS) and anticorrelated RDS (aRDS), in which corresponding dots between the two eyes have opposite contrast that generate false matches (Fig. 8A; Cumming and Parker, 1997). Stimulus presentation in the awake state allowed to evoke substantially stronger neuronal responses to RDS compared to the anesthetized state, activating a higher fraction of neurons and with larger calcium transients on average (data not shown), while preserving the overall disparity tuning of individual neurons across states (La Chioma et al., 2019). Many neurons across areas exhibited clear responses to cRDS and aRDS, with reliable activation by a limited range of disparities (Fig. 8B). Compared to dichoptic gratings as measured in anesthetized mice, cRDS presented to awake mice activated a much higher proportion of cells (percentage of cells responsive to cRDS, mean ± SEM across planes, V1, 47.2% ± 3.6%; LM, 44.4% ± 6.6%; RL, 32.9% ± 2.5%). When comparing the fraction of responsive cells between experiments with grating and RDS stimuli, it should be borne in mind that ROI segmentation was performed with two different methods, namely manual segmentation for gratings and automated segmentation for RDS (see Materials and Methods). On average, 48.1%, 78.7%, and 55.7% of responsive neurons in areas V1, LM, and RL, respectively, were disparity-tuned to cRDS (see Materials and Methods). In contrast, aRDS stimuli activated far fewer neurons (V1: 15.7%, LM: 7.6%, RL: 13.6%). Across areas, 3–4% of all responsive cells were disparity-tuned to both cRDS and aRDS (Fig. 8C). Notably, LM showed a significantly higher proportion of cells tuned only to cRDS compared to V1 and RL, while having a smaller proportion of cells tuned only to aRDS (Fig. 8C). Using more stringent criteria for defining responsive cells did not qualitatively change the proportions of disparity-tuned cells (data not shown).

**Figure 8.**
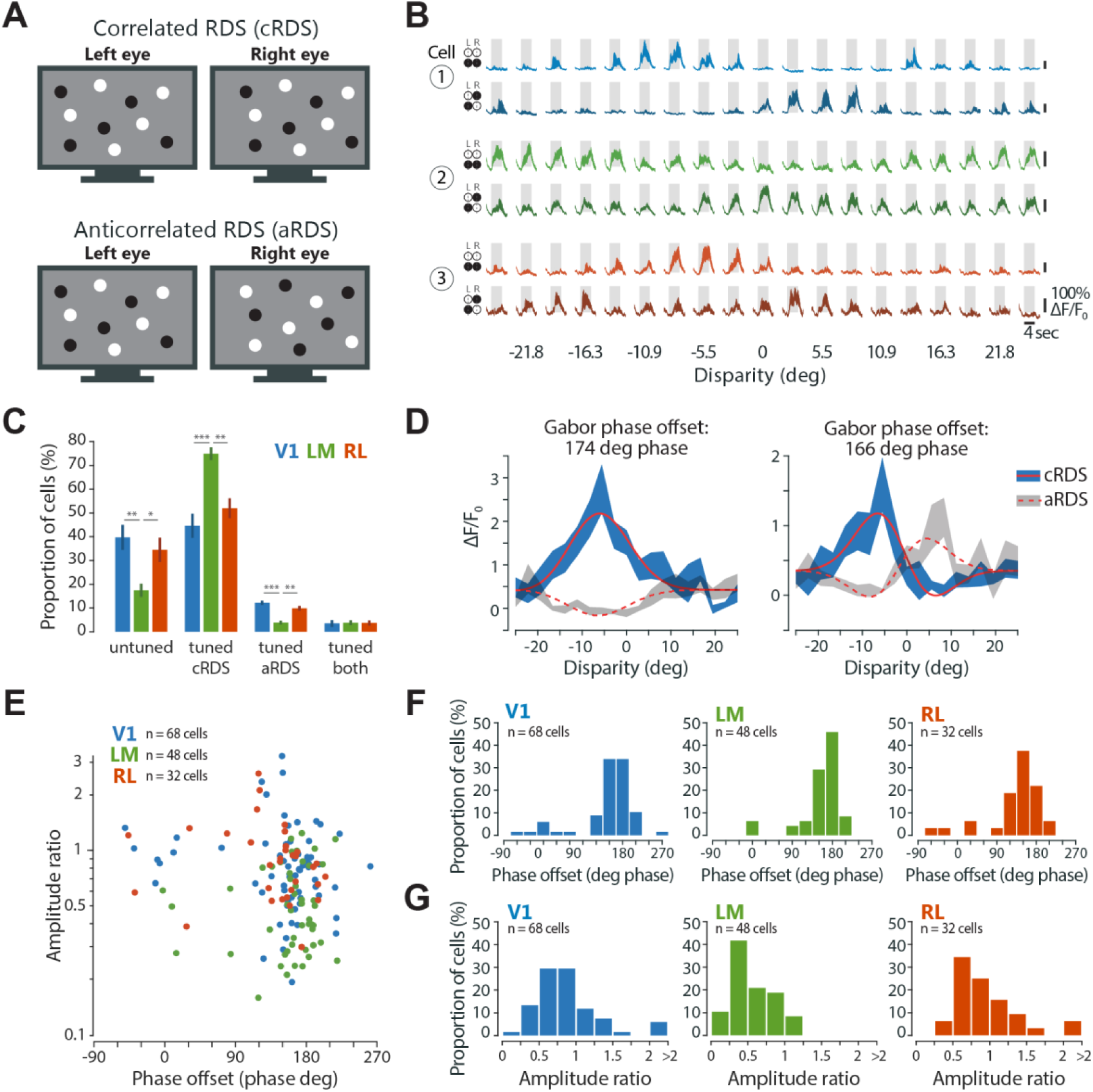
Area-specific responses to anticorrelated random dot stereograms. (A) Schematic of correlated and anticorrelated random dot stereogram stimuli. (B) Visually evoked calcium traces (ΔF/F_0_) of three example neurons, one from each area, in response to cRDS (upper traces, lighter shading) and aRDS (lower traces, darker shading). Fluorescence time courses are plotted as mean ΔF/F_0_ ± SEM (shaded areas) calculated across stimulus trials (10 repeats). Gray boxes, duration of stimulus presentation (4 s), bottom edge indicates baseline level (0% ΔF/F_0_). (C) Percentage of disparity-tuned and untuned neurons, mean ± SEM across imaging planes. Statistical tests: Kruskal-Wallis test across areas for each tuning group separately, followed by Bonferroni-corrected Mann-Whitney tests, with asterisks denoting significance values of post hoc tests. (D) Tuning curve fit with Gabor functions for two example neurons in response to cRDS (solid lines) and aRDS (dashed lines). Note tuning inversion between cRDS and aRDS, as indicated by a Gabor phase offset of around ±180 deg phase. (E) Gabor amplitude ratio plotted against Gabor phase offset between cRDS and aRDS tuning curves, for individual neurons tuned to both stimuli (V1, n = 68 cells, 9 planes, 4 mice; LM, n = 48 cells, 8 planes, 3 mice; RL, n = 32 cells, 9 planes, 4 mice). (F) Distributions of Gabor phase offset between cRDS and aRDS tuning curves for each area. (G) Distributions of Gabor amplitude ratio between cRDS and aRDS tuning curves for each area.

Neurons tuned to both cRDS and aRDS often exhibited a tuning inversion: disparities evoking strong responses with correlated stimuli caused weak activations with anticorrelated stimuli and vice versa (Fig. 8B). The tuning inversion indicates that these neurons operate only as local disparity detectors, disregarding contrast sign and the global consistency of image features between the two eyes, thereby resulting in responses to false matches (Cumming and Parker, 1997). To quantitatively describe the relation between correlated and anticorrelated responses, we fitted the disparity tuning curves of each neuron with Gabor functions (Fig. 8D; Cumming and Parker, 1997; see Materials and Methods for details). The offset in Gabor phase between cRDS and aRDS tuning curves quantifies the degree of tuning inversion, with values of 180 deg phase indicating full inversion and values around 0 deg phase indicating no change. Likewise, the ratio of the Gabor amplitude between aRDS and cRDS tuning curves quantifies the change in disparity sensitivity, with values below 1 signifying a reduced response modulation to aRDS relative to cRDS. Plotting these two measures against each other reveals that most cells across areas underwent tuning inversion (phase offset of around 180 deg phase) and a decrease in modulation to aRDS (amplitude ratio < 1, Fig. 8E). While the incidence of tuning inversion is similar across all three areas (Fig. 8F; circular non parametric multi-sample test for equal medians, across cells, p = 0.1889), area LM shows a significantly lower amplitude ratio (Fig. 8G; Kruskal-Wallis test across cells, Χ^2^ = 26.987, p = 1.3800– 06; post hoc Bonferroni-corrected Mann-Whitney tests, V1 versus LM, p = 1.493e–05; V1 versus RL, unadjusted p = 0.5372; LM versus RL, p = 3.981e–05; sign test, left-tailed for median <1, across cells: V1, p = 6.541e–05; LM, p = 7.569e–10; RL, p = 0.0551). Thus, area LM features a smaller fraction of neurons tuned to aRDS, and these neurons display weaker disparity modulation to aRDS, compared to V1 and RL. We conclude that mouse area LM performs a higher-lever analysis of disparity signals compared to V1 and RL.

## Discussion

Our study shows that the integration of signals from both eyes is a prominent feature of mouse visual cortex. Large fractions of neurons in areas V1, LM, and RL, even when classified as monocular by conventional ocular dominance measurements, are in fact binocular, in the sense that their activity can be strongly facilitated or suppressed by simultaneous input from both eyes and over a specific range of interocular disparities. We observed some degree of fine-scale spatial organization for disparity tuning, as neurons with similar disparity preference are clustered within a horizontal range of ∼10 μm. Moreover, similarly tuned neurons have higher noise correlations, suggesting that they are more strongly interconnected or share common input. Comparing responses to binocularly correlated and anticorrelated RDS, we found that area LM shows a higher selectivity for correlated stimuli, while being less sensitive to anticorrelated RDS.

### Disparity processing is widespread across mouse visual areas V1, LM, and RL

The overall high disparity selectivity found across mouse visual areas V1, LM, and RL, which harbor the largest, continuous representation of the binocular visual field in the visual cortex, matches the widespread distribution of disparity processing throughout most of the visual cortex of carnivorans and primates. The binocular disparity signals that these neurons carry are potentially critical for depth perception, because they are essential for the construction of stereopsis by the visual system. Thus, the abundance of disparity signals found in all three areas strongly suggests that mice do use binocular disparity as a depth cue to estimate object distances. While it is not fully understood how rodents move their two eyes to perceive the environment and enable binocular vision (Wallace et al., 2013; Meyer et al., 2018, 2020; Michaiel et al., 2020), mice are capable of stereoscopic depth perception, sharing at least some of the fundamental characteristics of stereopsis with carnivorans and primates (Samonds et al., 2019).

Studies in these species have long sought to identify a cortical region specifically dedicated to stereoscopic depth processing, but failed in achieving this goal (Parker, 2007). For other visual object features, specific cortical regions have been shown to be particularly relevant. For example, primate areas V4 and MT are considered crucial centers for color and motion perception, respectively (Lueck et al., 1989; Born and Bradley, 2005). Why is binocular disparity processing so widespread across multiple areas? One possibility is that disparity processing relies on the concomitant recruitment of several areas. Similar disparity signals generated in these areas might then be differentially combined with information about other aspects of visual stimuli, such as motion, contrast, and shape, or with information deriving from other sensory modalities, in order to construct the percept of a 3D object. Another possibility is that different areas do form specialized representations of binocular disparity, thereby playing distinct disparity-related roles in constructing a 3D percept (Roe et al., 2007), such as encoding near or far space (Nasr and Tootell, 2018; La Chioma et al., 2019), supporting reaching movements, or computing visual object motion across depth (Czuba et al., 2014; Sanada and DeAngelis, 2014).

### Asymmetric disparity tuning curves

The disparity tuning width of individual neurons as measured with dichoptic gratings was on average similar across areas V1, LM, and RL. Most neurons in these areas showed symmetric tuning. A small subset of cells (∼15%) in areas V1 and LM, exhibited an asymmetric tuning curve (Fig. 5D). Most of these cells had a disparity preference maximally different (180 deg phase) from the overall peak in the distribution of disparity preferences in each recording. A possible interpretation of the disparity tuning asymmetry of some cells might be related to the two proposed mechanisms underlying disparity selectivity: the position-shift and the phase-shift mechanism, which result in symmetric and asymmetric disparity tuning curves, respectively (Qian, 1997; Cumming and DeAngelis, 2001; Tsao et al., 2003). A disparity phase difference of 180 deg corresponds to 50 absolute degrees (a full grating cycle at 0.01 cpd corresponds to 100 absolute degrees). Assuming that the population peak disparity reflects the visual field location of retinal correspondence between the eyes (vertical meridian of zero retinal disparity), a position-shift mechanism alone would require a spatial offset between left and right eye RFs of 50 degrees. This value is unrealistically large, considering the typical RF size of cells in V1 and LM (average diameter of ∼25 and ∼40 deg, respectively; Smith and Häusser, 2010; Van den Bergh et al., 2010; Bonin et al., 2011; Vaiceliunaite et al., 2013; Smith et al., 2017), and that left and right eye RFs are at least partially overlapping in most V1 neurons (Sarnaik et al., 2014). It follows that cells in V1 and LM with a disparity preference of 0 deg phase might predominantly rely on a phase-shift mechanism, hence showing asymmetric tuning curves. By contrast, neurons in RL, possibly due to larger RFs compared to V1 and LM (de Vries et al., 2020), might rely less on a phase-shift mechanism.

### Responses to binocular correlation and anticorrelation across visual cortex

Generating coherent stereo depth perception requires the visual system to extract binocular disparity from the visual inputs to both eyes and match the image elements in one eye to the corresponding elements in the other eye. With aRDS, i.e. image elements having opposite contrast between the two eyes, the neurons’ RFs are presented with local matches that do not correspond to globally coherent matches. Human observers, indeed, perceive stereo depth with cRDS, but generally do not with aRDS (Julesz, 1971; Cogan et al., 1993, 1995; Cumming et al., 1998; Zhaoping and Ackermann, 2018). In primates, V1 neurons maintain disparity selectivity to aRDS despite an inversion of the tuning profile (Cumming and Parker, 1997). This indicates that V1 neurons perform a low-level analysis of disparity signals and that further disambiguation in downstream areas is necessary to discard false matches and generate stereo depth perception (Parker, 2007). Areas MT and MST in the primate dorsal stream show anticorrelated responses as strong as in V1 (Takemura et al., 2001; Krug, 2004). In contrast, in the ventral stream, response to binocular anticorrelation is reduced in area V4 (Tanabe et al., 2004) and is abolished further down the ventral pathway in the inferior temporal cortex (Janssen et al., 2003). These findings indicate that the ventral stream solves “the correspondence problem” by performing a higher-level analysis of disparity signals, which corresponds more closely to the perception of stereo depth (for review, see Verhoef et al., 2016). Together, these studies have helped establishing an areal hierarchy for depth processing in the primate visual system.

Accumulating evidence supports the view that also the rodent visual cortex is organized into two subnetworks, which may share similarities with the ventral and dorsal streams of primates. According to anatomical classifications based on corticocortical connectivity, higher visual areas appear to be subdivided into two distinct groups (Wang et al., 2011, 2012). In one group, areas LM, LI, POR, and P, located lateral, i.e. ventral, to V1, are more densely interconnected and preferentially target ventral regions of the cortex. In the other, dorsal group, areas AL, RL, A, AM, and PM, are located primarily anterior and medial to V1, and provide input predominantly to dorsal regions of the cortex (Wang et al., 2011, 2012). While supported by anatomical data, the existence of distinct processing streams in rodents is still quite speculative, as functional evidence to support this concept is presently scant compared to primates, and is mostly based on differences in tuning for select RF properties, like spatial and temporal frequency, orientation and motion direction (Andermann et al., 2011; Marshel et al., 2011; Roth et al., 2012; Tohmi et al., 2014; Murakami et al., 2017; Smith et al., 2017; for reviews, see Laramée and Boire, 2014; Glickfeld and Olsen, 2017).

In the present study, we found that the sensitivity to anticorrelated stimuli was similarly strong in areas V1 and RL, indicating that false matches are not fully rejected at this stage of disparity processing. In this respect, area RL, considered to be part of a putative dorsal subnetwork of mouse visual cortex, resembles areas MT and MST in the primate dorsal stream. In contrast, area LM showed a higher proportion of neurons tuned only to cRDS, while there were significantly fewer neurons tuned to aRDS, and their responses were weaker compared to V1 and RL. These findings suggest that area LM, which several studies consider part of a putative ventral subnetwork of mouse visual cortex, performs a higher-level analysis of disparity signals, reminiscent of V4 and inferior temporal cortex in the primate ventral stream.

In summary, our results show that sensitivity for binocular disparities is widespread across areas of mouse visual cortex, supporting the idea that mice might use this cue for depth perception (Samonds et al., 2019). Our findings also demonstrate areal specializations for disparity processing (La Chioma et al., 2019) that support a subdivision of mouse visual cortex into ventral and dorsal processing streams, which may share features with those in primates.

## Conflict of interest statement

The authors declare no competing interests.

## Acknowledgements

We thank Pieter Goltstein for setting up intrinsic signal imaging hardware and software. We are grateful to Max Sperling for the development and maintenance of data acquisition software. This work was supported by the Max Planck Society and a Boehringer Ingelheim Ph.D. fellowship to A.L.C.

## Materials and Methods

### Virus injection and cranial window implantation

All experimental procedures were carried out in accordance with the institutional guidelines of the Max Planck Society and the local government (Regierung von Oberbayern). A total of 13 female adult C57/BL6 mice were used, housed with littermates (3-4 per cage) in a 12:12 hr light-dark cycle in individually ventilated cages, with access to food and water *ad libitum*.

Cranial window implantations were performed at 10–12 weeks of age, following the procedures described in (La Chioma et al., 2019). Briefly, mice were anesthetized by intraperitoneal injection of a mixture of fentanyl (0.075 mg/kg), midazolam (7.5 mg/kg), and medetomidine (0.75 mg/kg). Virus injections were performed through a circular craniotomy (4-5 mm diameter) into the right hemisphere (images in Fig. 1C,E were mirrored for consistency with cited references), at 3–5 sites in the binocular region of V1 and ∼0.5–1 mm more lateral (corresponding to the location of areas LM and RL), using AAV2/1.Syn.mRuby2.GSG.P2A.GCaMP6s.WPRE.SV40 (Rose et al., 2016). Following injections, the craniotomy was sealed flush with the brain surface using a glass cover slip. A custom machined aluminum head-plate was attached to the skull using dental cement to allow head-fixation during imaging. Expression of the transgene was allowed for 2.5–3 weeks before imaging.

### Intrinsic signal imaging

Intrinsic signal imaging was used to localize areas V1, LM, and RL. Imaging was performed under anesthesia 2-4 weeks after cranial window implantation, as detailed in (La Chioma et al., 2019).

### In vivo two-photon imaging

Two-photon imaging was performed 3–17 weeks after cranial window implantation for experiments under anesthesia (8 mice total) and 6-12 weeks after cranial window implantation for experiments in awake animals (5 mice total). For imaging under anesthesia, mice were initially anesthetized by intraperitoneal injection of a mixture of Fentanyl (0.030 mg/kg), Midazolam (3.0 mg/kg), and Medetomidine (0.30 mg/kg). Additional anesthetic mixture (25% of the induction dose) was injected subcutaneously 60 min after the initial injection and then every 30-40 min to maintain anesthesia. Images were acquired using a custom-built two-photon microscope (La Chioma et al., 2019) equipped with an 8 kHz resonant galvanometer scanner, resulting in frame rates of 17.6 Hz at an image resolution of 75×900 pixels (330×420 μm). The illumination source was a Ti:Sapphire laser with a DeepSee pre-chirp unit (Spectra Physics MaiTai eHP), set to an excitation wavelength of 940 nm. Laser power was 10–35 mW as measured after the objective (16x, 0.8 NA, Nikon). For awake imaging, the animals were head-fixed on top of an air suspended Styrofoam ball (diameter 20 cm), allowing the mouse to run during stimulus presentation and data acquisition (Dombeck et al., 2007).

### Monitoring eye position

During two-photon imaging, both eyes were continuously imaged with an infrared video camera (The Imaging Source, frame rate 30 Hz). Pupil position and diameter were monitored online using custom-written software (LabVIEW, National Instruments) based on (Sakatani and Isa, 2007). Analysis of pupil position was also performed post hoc to test whether either eye had changed position over the course of the experiment. Approximately 10% of the imaging experiments under anesthesia were discarded owing to eye drifts.

### Visual stimulation: Dichoptic stimulation

All visual stimuli presented during two-photon imaging in anesthetized mice (Figs. 1 to 7) were displayed through a haploscope, consisting of two separate mirrors and two separate display monitors to enable independent stimulation of each eye (see Fig. 1D). Each mirror (silver coated, 25 × 36 × 1.05 mm, custom-made, Thorlabs), mounted on a custom designed, 3D printed plastic holder, was independently positioned at an angle of approximately 30 deg to the longitudinal axis of the mouse, contacting the snout 2-4 mm anterior to the medial palpebral commissure of each eye. A shield made of black paper board and tape was used to prevent stimulus cross-talk between eyes and monitors. Each mirror redirected the field of view of each eye onto a separate display monitor located on each side of the animal at a distance of 21 cm, with an actual stimulation area subtending 65 deg in elevation and 70 deg in azimuth for each eye. The two 21-inch LCD monitors (gamma corrected, refresh rate of 60 Hz, spatial resolution of 1600×900 pixels) were mounted on custom machined metal holders that allowed flexible and reproducible positioning of each monitor independently. To minimize light contamination of data images from visual stimulation, the LED backlight of the monitors were flickered at 16 kHz such that they were synchronized to the line clock of the resonant scanner (Leinweber et al., 2014). As a result, the LED backlight was only active during the turnaround intervals of the scan phase, which were not used for image generation (mean luminance with 16 kHz flickering: white, 5.2 cd/m^2^; black 0.01 cd/m^2^).

For dichoptic stimulation with RDS during two-photon imaging in awake mice (Fig. 8), eye shutter glasses (3D Vision 2, Nvidia) were used (La Chioma et al., 2019). The glasses consisted of a pair of liquid crystal shutters, one for each eye, that rapidly (60 Hz) alternated their electro-optical state – i.e. either occluded or transparent to light. In one frame sequence, the left eye shutter was occluded while the right eye shutter was transparent, and vice versa for the next frame, with alternations synchronized to the monitor refresh rate (120 Hz). Synchrony between the shutter glasses and the monitor was accomplished with an infra-red wireless emitter. The display monitor (Acer GN246HL, 24 inches, 120 Hz refresh rate, spatial resolution 1600×900 pixels) was placed in front of the mouse at a distance of 13 cm from the eyes (luminance measured through the transparent shutter: white, 21.6 cd/m^2^; black 0.05 cd/m^2^). To reduce light contamination of two-photon images from visual stimulation, the microscope objective was shielded using black tape. Visual stimuli were generated using custom-written code for Matlab (MathWorks) with the Psychophysics Toolbox (Brainard, 1997; Kleiner et al., 2007). For dichoptic stimulation through the haploscope, the code was run on a Dell PC (T7300) equipped with a Nvidia Quadro K600 graphics card and using a Linux operating system to ensure better performance and timing in dual-display mode, as recommended (Kleiner, 2010). For dichoptic stimulation through eye shutter glasses, the code was run on a Dell PC (Precision T7500) equipped with a Nvidia Quadro K4000 graphics card and using Windows 10.

### Visual stimulation: Retinotopic mapping

When using the haploscope for dichoptic stimulation (Figs. 1 to 7), retinotopic mapping was performed to ensure that the field of view of each eye was roughly redirected onto the central area of its display monitor. Visual stimuli consisted of vertical and horizontal patches of drifting gratings, presented to each eye in 6 vertical (size 16×65 deg, w × h) and 5 horizontal (size 70×13 deg, w × h) locations (eight consecutive directions in pseudorandom sequence; SF, 0.05 cpd; TF, 2 Hz; duration of each stimulus patch, 4 sec; inter-stimulus interval, 2 sec; number of stimulus trials, 2–4). Retinotopic maps for each eye were generated immediately after completing stimulation (see below for details on analysis). If the center of the ensemble receptive field (RF) of one eye was closer than approximately 20 deg to the screen edge, the position of the haploscope mirror of that eye was adjusted and the retinotopic mapping was repeated.

### Visual stimulation: Monocular drifting gratings

When using the haploscope for dichoptic stimulation (Figs. 1 to 7), slight misalignment of the two mirrors could cause the artefactual rotation of one eye’s field of view relative to that of the other eye. To estimate the rotation offset between each eye’s field of view (see below), monocular drifting gratings were used. Sinusoidal oriented gratings were presented, to each eye separately, at 12 or 16 equally spaced drifting directions (30–360 deg or 22.5–360 deg). Gratings were displayed in full-field and at 100% contrast, at a SF of 0.05 cpd and a TF of 2 Hz. Each stimulus was displayed for 3 sec (6 grating cycles, randomized initial spatial phase) preceded by an inter-stimulus interval of 2 sec with a blank (gray) screen with the same mean luminance as during the stimulus period. During presentation of a grating stimulus on one monitor, the other monitor displayed a blank screen. Each stimulus was repeated for 2–3 trials, in pseudorandomized sequence across drifting directions and eyes. In a given experiment, all subsequent stimuli presented through the haploscope were displayed by correcting for the eye rotation offset, which was estimated by online analysis of the neuronal responses, assuming matched orientation preference between the two eyes in adult mice (Wang et al., 2010; see below for details on the analysis).

To measure OD as well as facilitation/suppression (Fig. 3), sinusoidal oriented gratings were presented with the following stimulus parameters: two directions of drifting vertical gratings (90 deg, rightward; 270 deg, leftward); one of three possible SFs, spaced by 2 octaves, among 0.01 cpd, 0.05 and 0.10 cpd; TF of 2 Hz; stimulus interval of 2 sec; inter-stimulus interval of 4 sec; each stimulus was repeated for 6 trials, in pseudorandomized sequence across drifting directions and eyes, and interleaved with dichoptic drifting gratings as a part of the same stimulation block.

### Visual stimulation: Dichoptic drifting gratings

Sinusoidal, drifting vertical gratings were dichoptically presented to both eyes, at varying interocular disparities. Different interocular grating disparities were generated by varying the initial phase (position) of the grating presented to one eye relative to the phase of the grating presented to the other eye across the full grating cycle. Eight equally spaced phase disparities (45–360 deg phase, spacing 45 deg phase) were used at a SF of 0.01 cpd. For each stimulus, drift direction (leftward or rightward), TF, and SF were kept constant across eyes. The other grating stimulus parameters were the same as for the monocular drifting gratings. Gratings were presented in pseudorandomized sequence across disparities and drifting directions, with 5-6 trials for each stimulus condition.

### Visual stimulation: Random dot stereograms

Random dot stereograms (RDS) consisted of a pattern of random dots, presented to both eyes in a dichoptic fashion. Between the left and the right eye stimulus patterns, a spatial offset along the horizontal axis was introduced to generate interocular disparities. A total of 19 different RDS conditions were presented, covering a range of disparities between –26.39 deg and +26.39 deg. The different RDS conditions were obtained by dividing the entire range of disparities into 19 nonoverlapping bins (bin width 2.78 deg) and assigning each bin to one RDS condition (e.g. [–1.39 +1.39], [+1.39 +4.17], etc.). Each RDS stimulus was presented for 4 sec, during which a new random pattern of dots was displayed every 0.15 sec. In each pattern, all dots had the same interocular disparity, randomly chosen within the 2.78 deg bin of that particular RDS condition; 50% of the dots (diameter 12 deg) were bright (brightness 77%) and 50% of the dots were dark (brightness 23%) against a gray background, with an overall density of 25%. For binocularly correlated RDS (cRDS), the dots were displayed with the same contrast between the two eyes, whereas for anticorrelated RDS (aRDS) the dots were displayed with opposite contrast between the two eyes. Each RDS condition was presented for 10 stimulus trials in pseudorandomized sequence, with individual trials separated by an inter-stimulus interval of 2 sec. RDS were displayed applying spherical correction for stimulating in spherical visual coordinates using a flat monitor.

### Analysis of imaging data

Imaging data were processed using custom-written Matlab software, as detailed in (La Chioma et al., 2019). Briefly, (1) images were aligned using a rigid motion registration; (2) regions of interest (ROIs) were selected by manually drawing circular shapes around cell somas; (3) the fluorescence time course of each cell was corrected for neuropil contamination and computed as F_cell_corrected_ = F_cell_raw_ *- r* x F_neuropil_, where F_cell_raw_ is the raw fluorescence time course of the cell extracted by averaging all pixels within the somatic ROI, F_neuropil_ was extracted from an annular neuropil ROI centered around the somatic ROI (3–13 μm from the border of the somatic ROI), and *r* is a contamination factor set to 0.7 (Kerlin et al., 2010; Chen et al., 2013). For experiments in awake mice (Fig. 8), imaging data were processed using the Suite2P toolbox in Matlab (Pachitariu et al., 2016), which entailed image registration, segmentation of ROIs, and extraction of calcium fluorescence time courses. Relative changes in fluorescence signals (ΔF*/* F_0_) were calculated, for each stimulus trial independently, as (F *-* F_0_)*/*F_0_, where F_0_ was the average over a baseline period of 1 sec immediately before onset of the visual stimulus.

### Analysis of retinotopic mapping of higher visual areas

Retinotopic maps for azimuth and elevation were generated using the temporal phase method (Kalatsky and Stryker, 2003) on images obtained with intrinsic signal imaging, as described (La Chioma et al., 2019). The boundary between V1 on the medial side and areas LM, AL, and RL on the lateral side was identified by a reversal at the vertical meridian, as indicated by the longer axis of the elliptically shaped contour on the vertical meridian. The boundaries between LM and AL, and between AL and RL were identified as a reversal near the horizontal meridian (Kalatsky and Stryker, 2003; Marshel et al., 2011; Garrett et al., 2014). The binocular regions of areas V1, LM, and RL were then specifically targeted for two-photon imaging, by using the blood vessels as landmarks, which could be reliably recognized in the two-photon images.

### Online analysis of retinotopic mapping

Data images of the recording were analyzed online (“on-the-fly”) pixel by pixel, by calculating a ΔF*/*F for each pixel and grating patch location, averaged across trials, for each eye independently. Azimuth and elevation maps were generated for each eye by counting, for each vertical and horizontal stimulus location presented to one eye, the number of pixels that best responded to it (only pixels with an averaged ΔF*/*F_0_ above zero for any given location were considered). Given the high number of total pixels in the images (675k), analyzed regardless of individual cell’s ownership, this online analysis provided a good estimate of the center of the overall RF across cells from that imaging plane, as confirmed by comparison to individual cells’ RFs analyzed *post hoc* (data not shown).

### Eye rotation offset online analysis

To estimate the eye rotation offset potentially caused by the haploscope mirrors (see above), data images containing responses to drifting gratings were analyzed online (“on-the-fly”) pixel by pixel, by calculating ΔF*/*F for each pixel and grating direction, averaged across trials. For each eye independently, responsive and orientation-tuned pixels were selected on the basis of an orientation selectivity index (OSI), scaled by the maximum relative fluorescence change, with OSI calculated for each pixel as the normalized length of the mean response vector (Ringach et al., 2002; Mazurek et al., 2014): 

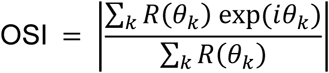

where *R*(Θ*k*) is the mean ΔF*/*F response to the orientation angle Θ*k*. The angle of the same mean response vector was taken as the preferred orientation of that pixel. For pixels that were selected for both eye-specific stimuli, the difference in preferred orientation between left and right eye was calculated and the average across all selected pixels was taken as the rotational offset between the eyes’ fields of view (dO). Subsequent stimulations (dichoptic gratings) were presented by correcting stimulus orientation by -dO/2 and +dO/2 for stimuli presented to the left and right eye, respectively.

### Responsive cells

Cells were defined visually responsive when ΔF_peak_*/*F_0_ *>* 4 x *σ*_baseline_ in at least 50% of the trials of the same stimulus condition, where ΔF_peak_ is the peak ΔF*/*F_0_ during the stimulus period of each trial, and *σ*_baseline_ is the standard deviation calculated across the F_0_ of all stimulus trials and conditions of the recording. For grating stimuli, the mean ΔF*/*F_0_ over the entire stimulus interval (2 sec) of each trial was calculated. For RDS stimuli, the mean ΔF*/*F_0_ of each trial was calculated over a time window of 1 sec centered around ΔF_peak_.

### Disparity selectivity index

For each cell responsive to dichoptic gratings, a disparity selectivity index (DI) was calculated, given by the normalized length of the mean response vector across the eight phase disparities of the drift direction that elicited the stronger activation (Scholl et al., 2013, 2015): 

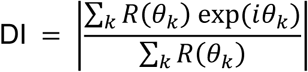

where *R*(Θ_*k*_) is the mean ΔF*/*F response to the interocular phase disparity Θ_*k*_. Cells were defined disparity-tuned to dichoptic gratings if DI>0.3. Using more stringent criteria for defining disparity-tuned cells (DI>0.5) did not qualitatively affect the results and the statistical significance of all analyses. In addition, using more stringent criteria for defining responsive cells (ΔF_peak_*/*F_0_ *>* 8 x σ_baseline_) did not result in a significant change in the DI distribution for each area (data not shown), ruling out that neuronal calcium signals with low signal-to-noise ratio affected the measurement of disparity selectivity with gratings. Note that the calculation of DI is based on a circular metric. As such, DI could be computed only for responses to dichoptic gratings, but not for responses to RDS, which are not circular. Cells were defined disparity-tuned to cRDS or aRDS when at least 50% of the tuning curve variance (*R*^*2*^) could be accounted for by the model fit (see below). Using more stringent criteria for defining cells responsive to cRDS or aRDS (ΔF_peak_*/*F_0_ *>* 8 x σ_baseline_) did not qualitatively affect the results and the statistical significance of all analyses (data not shown), suggesting that the measurement of disparity sensitivity to cRDS and aRDS across areas was not affected by low signal-to-noise ratios.

### *Ocular dominance inde*x

Ocular dominance was determined for responsive cells by calculating the ocular dominance index (ODI) using eye-specific responses to drifting gratings: 

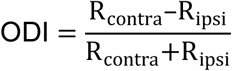

where R_contra_ and R_ipsi_ are the mean ΔF*/*F_0_ responses (across trials) to the preferred grating direction and SF presented to either the contra- or ipsilateral eye, respectively. Contralateral and ipsilateral dominance are indicated by an ODI of 1 or −1, respectively. A cell equally activated by either eye stimulation has an ODI = 0.

### *Facilitation and suppression inde*x*es*

To quantify the response facilitation or suppression at the preferred and least-preferred disparity, respectively, a facilitation index (FI) and a suppression index (SI) were computed for every neuron responsive to both dichoptic and monocular gratings, at the neuron’s preferred SF, defined as: 

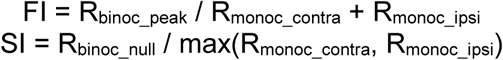

R_binoc_peak_ and R_binoc_null_ are, respectively, the largest and smallest response evoked among the eight disparities; R_monoc_contra_ and R_monoc_ipsi_ are the responses to monocular gratings presented, respectively, to either the contra- or ipsilateral eye. FI values above 1 indicate a facilitatory interaction of the eye-specific inputs, while SI values below 1 correspond to suppressive binocular interactions. Moreover, FI values above 1 and SI values below 1 in principle indicate a non-linear integration mechanism through which responses are facilitated and suppressed, respectively. It should be borne in mind, however, that the responses measured in this study consist of the visually-evoked fluorescence signal of GCaMP6s, which provides an indirect measure of the spiking activity of neurons (Hendel et al., 2008; Grienberger and Konnerth, 2012; Lütcke et al., 2013; Rose et al., 2014). Owing to the non-linear relationship between action potential firing and the GCaMP6 fluorescence signal, only the presence of facilitation or suppression can be reported, without inferring the precise linear/non-linear nature of the binocular interaction shown by a cell.

### Disparity tuning curve fit

Disparity tuning curves obtained with dichoptic gratings were fitted with an asymmetric Gaussian function using single trial responses (Hinkle and Connor, 2005), as follows: 

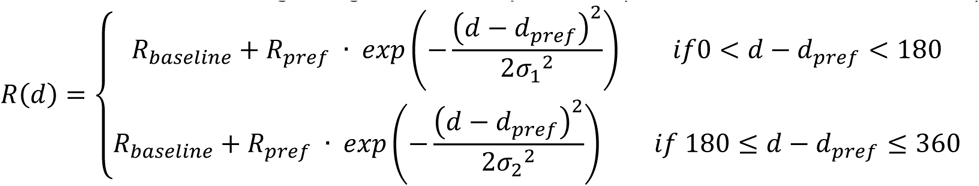

where *R*_*baseline*_ is the baseline response, *R*_*pref*_ is the response to the preferred disparity, *σ*_*1*_ and *σ*_*2*_ are the tuning width parameters for the left and right sides, respectively. The tuning curve asymmetry as plotted in Figure 5D was quantified as *σ*_*2*_ - *σ*_*1*_. Disparity preferences of responsive cells for dichoptic gratings were given by the fit parameter *d*_*pref*_when at least 50% of the tuning curve variance (*R*^*2*^) could be accounted for by the model fit.

Disparity tuning curves obtained with cRDS or aRDS were fitted with a Gabor function using single trial responses, as follows: 

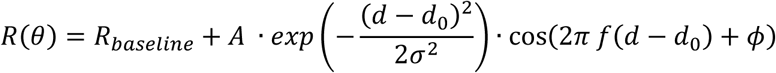

where *A* is the amplitude, *d*_*0*_ is the Gaussian center, *σ* is the Gaussian width, *f* is the Gabor frequency, and *ϕ*is the Gabor phase. When cells were responsive to both cRDS and aRDS stimuli, the tuning curves obtained with cRDS and aRDS were fitted using the same values of *R*_*baseline*_, *d*_0_, *σ*, and *f*, for both curves, but different values for *A* and *ϕ*(*A*_*c*_, *A*_*a*_, *ϕ*_*c*_, *ϕ*_*a*_, with the subscripts *c* and *a* referring to the tuning curve obtained with cRDS and aRDS, respectively; Cumming and Parker, 1997).

### Noise correlations

Noise correlations were calculated between all possible pairs of disparity-tuned neurons from the same imaging plane. The single-trial ΔF*/*F responses of a given cell to dichoptic gratings were Z-scored with respect to the mean across trials. Pairwise noise correlations were then computed using the Pearson’s linear correlation coefficient. While fluorescence signals were corrected for contamination by surrounding neuropil and neighboring cells, noise correlations might still be affected by potentially residual neuropil contamination. For this reason, only pairs separated by at least 20 μm were considered.

### Population decoding with SVM

To estimate how much information about binocular disparity is carried by the joint activity of populations of neurons in each area, a population decoding approach based on support vector machines (SVM; Cortes and Vapnik, 1995) was employed. The decoding approach was designed to estimate which disparity, among all eight possible grating disparities, was actually presented. This discrimination among eight distinct classes was redefined as a series of binary classifications (“multi-class classification”), in which SVM were used to find the hyperplane that best separated neuronal activity data points of one class (grating disparity) from those of another class (“binary classification”). The accuracy in classifying data points of the two stimulus conditions with increasing numbers of neurons was evaluated.

The dataset consisted of pseudo-populations generated by pooling neurons, from a given area and responsive to dichoptic gratings, from all individual recordings across animals. To take into account that distributions of disparity preferences had a population peak disparity that was characteristic for each individual recording (see Fig. 4A), the tuning curves of neurons from a given recording were shifted by dP deg phase, where dP is the difference between the population peak disparity characteristic of that recording and 180 deg phase. For each neuron in the dataset, the ΔF*/*F_0_ response of each trial was split in b = 6 bins of 0.5 sec, including 4 bins during the stimulus period (2 sec) and 2 bins immediately following it, and the mean ΔF*/*F_0_ of each bin was taken as one activity data point. As such, for each neuron and disparity, there were ap activity points, with ap = t x b, where t = 6 trials for each disparity and b = 6 bins.

For each decoding session, a subpopulation of N neurons was randomly sampled from the pseudo-population. A matrix of data points was constructed, with N columns (neurons, corresponding to the “features”) and ap x d rows (activity points x disparities, corresponding to the “observations”). The data matrix was divided into two separate sets, a training set and a test set. The training set included 0.9 x ap randomly chosen activity points for each disparity; the test set included the remaining 0.1 x ap activity points (“10-fold cross-validation”). A multi-class decoder was constructed by training 28 distinct binary classifiers, each considering only two different disparities as the two classes, and exhausting all combinations of disparity pairs (“one-vs-one”). The identity of each observation of the training set was also provided to every classifier (“supervised classification”). Then, the multi-class decoder was probed on the test set. Each observation of the test set was evaluated by each of the 28 binary classifiers to predict its class (disparity). The class identity that was more frequently predicted across the 28 classifications was taken as the predicted class identity of that observation. This evaluation was performed for every observation of the test set. The procedure was then repeated on a different training set and test set, across all 10 folds, to produce an average accuracy estimate of the decoder for a given subpopulation of N neurons. 20 different random resamplings of N neurons from the pseudo-population were performed and the outcomes were averaged to generate a measure of decoding performance of a given N, as reported in Figure 7. Significance levels for classification accuracy were determined by using a similar decoding procedure but training decoders on training sets in which the identities of the observations were randomly shuffled, repeating the shuffling 100 times. The binary classifiers consisted of SVM with a linear kernel. The decoding procedures were performed using custom-written routines based on the function *fitecoc* (kernel scale, 1; box constraint, 1) as part of the Statistics and Machine Learning Toolbox in Matlab (Mathworks).

### Statistical analyses

All data and statistical analyses were performed using custom-written Matlab code (MathWorks). Sample sizes were not estimated in advance. No randomization or blinding was performed during experiments or data analysis. Data are reported as mean with standard error of the mean (mean *±* SEM). Data groups were tested for normality using the Shapiro-Wilk test in combination with a skewness test and visual assessment (Ghasemi and Zahediasl, 2012). Comparisons between data groups where made using the appropriate tests: one-way ANOVA, Kruskal-Wallis test, circular non parametric multi-sample test for equal medians (Fig. 8F; Berens, 2009). For multiple comparisons, Bonferroni correction was used. One-sample, one-sided t-test for mean > 0 was used in Figure 6A; one-sample, one-sided sign test for median < 1 was used in Figure 8G; all other tests were two-sided. Significance levels for spatial clustering in Figure 4B were determined by permutation tests. In each permutation (n = 1000 permutations), the xy positions of all neurons from a given area were randomly shuffled and the mean (across all disparity-tuned neurons of a given area) difference in disparity preference for each distance bin was calculated; 95% confidence intervals were computed as 2.5–97.5 percentiles of all permutations. Significance levels for noise correlations in Figure 6C,D were determined by permutation tests. In each permutation (n = 1000 permutations), noise correlations of all pairs of disparity-tuned neurons from each imaging plane of a given area were randomly shuffled and the mean (across planes) noise correlations for each bin of distance (Fig. 6C) or bin of difference in disparity preference (Fig. 6D) were calculated; 95% confidence intervals were computed as 2.5–97.5 percentiles of all permutations. The statistical significances are reported in the Figures, with asterisks denoting significance values as follows: * p < 0.05, ** p < 0.01, *** p < 0.001.

### Data and code availability

Data and analysis code used in this study are stored and curated at the Max Planck Computing and Data Facility Garching (Munich, Germany) and are available from the corresponding authors upon reasonable request.

## Notes

### Competing Interest Statement

The authors have declared no competing interest.

